# spongEffects: ceRNA modules offer patient-specific insights into the miRNA regulatory landscape

**DOI:** 10.1101/2022.03.29.486212

**Authors:** Fabio Boniolo, Markus Hoffmann, Norman Roggendorf, Bahar Tercan, Jan Baumbach, Mauro A. A. Castro, A. Gordon Robertson, Dieter Saur, Markus List

## Abstract

**Motivation:** Cancer is one of the leading causes of death worldwide. Despite significant improvements in prevention and treatment, mortality remains high for many cancer types. Hence, innovative methods that use molecular data to stratify patients and identify biomarkers are needed. Promising biomarkers can also be inferred from competing endogenous RNA (ceRNA) networks that capture the gene-miRNA gene regulatory landscape. Thus far, the role of these biomarkers could only be studied globally but not in a sample-specific manner. To mitigate this, we introduce spongEffects, a novel method that infers subnetworks (or modules) from ceRNA networks and calculates patient- or sample-specific scores related to their regulatory activity.

**Results:** We show how spongEffects can be used for downstream interpretation and machine learning tasks such as tumor classification and for identifying subtype-specific regulatory interactions. In a concrete example of breast cancer subtype classification, we prioritize modules impacting the biology of the different subtypes. In summary, spongEffects prioritizes ceRNA modules as biomarkers and offers insights into the miRNA regulatory landscape. Notably, these module scores can be inferred from gene expression data alone and can thus be applied to cohorts where miRNA expression information is lacking.

**Availability:** https://bioconductor.org/packages/devel/bioc/html/SPONGE.html

**Supplementary information:** Supplementary data are available at Bioinformatics online.

## INTRODUCTION

Despite recent advances in screening, diagnosis, and prognosis, and an increased understanding of the mechanisms driving tumorigenesis, progression, and maintenance, cancer deaths are estimated to be ~400,000/year by 2040 in the US alone (Rahib *et al*., 2021), highlighting the need for innovative approaches for the diagnosis, monitoring, and treatment. In addition to the traditional study of genome variation and gene expression, microRNAs (miRNAs) have been assessed as potential biomarkers, given their important roles in regulating gene expression both in physiological conditions and in cancer (Liu *et al*., 2018). Mature miRNAs are short non-coding RNAs with lengths of ~20-23 nucleotides that play an important role in regulating gene product abundance (Kartha and Subramanian, 2014). miRNAs are involved in regulating at least half the genes in the human genome (Friedman *et al*., 2009) and are dysregulated in many diseases (Jiang *et al*., 2009). They function via the targeting of RNAs that will be degraded or whose translation will be impeded (Bartel, 2009). miRNA-target recognition is facilitated via matching the seed region at the 5’ end of the miRNA with the 3’ binding site of the target (canonical targeting) or via the action of additional regions outside of the seed that can contribute to the recognition of the target (non-canonical targeting) (McGeary *et al*., 2019). Salmena et al. proposed the competitive endogenous RNA (ceRNA) hypothesis (Salmena *et al*., 2011), which suggests that RNAs with miRNA binding sites compete for a limited pool of miRNAs, giving rise to a complex gene regulatory network (Tsang *et al*., 2010). Past studies have identified genes that act as ceRNAs (Salmena *et al*., 2011; Tay *et al*., 2014) and have associated them with different RNA classes, like messenger RNAs (mRNAs), circular RNAs, pseudogenes, transcripts of 3’ untranslated regions (UTRs), and long non-coding RNAs (lncRNAs) (Dykes and Emanueli, 2017). In coding genes, effective miRNA targeting (i.e., reducing transcript levels) largely occurs in 3’UTRs (Agarwal *et al*., 2015). lncRNAs have recently received particular attention in the framework of ceRNAs, suggesting the key role these molecules may play in binding miRNAs and indirectly regulating the expression of protein-coding genes sharing the same miRNA binding site region (Wang *et al*., 2022). It is important to note that the key actors in these modules are not the miRNAs but rather the ceRNAs that influence the expression of other ceRNAs by using miRNAs as a limited resource.

A range of computational methods has been developed to identify potential miRNA-target interactions based on different inputs, such as gene expression data, protein-protein interaction networks, and sequence information (Muniategui *et al*., 2013). Such methods often result in a large number of inferred interactions that, in return, suggest complex ceRNA networks that are typically difficult to explore. The key problem is to dissect these networks into functional units, or modules, that could play a role in specific tissues or diseases (Choobdar *et al*., 2019). The identification of ceRNA modules could highlight the biological relevance of miRNA regulation in specific processes while potentially identifying new prognostic biomarkers and therapeutic targets for clinical applications.

While tools for inferring ceRNA networks exist (Zhang *et al*., 2022), it is not straightforward to identify functional modules with biological or clinical relevance. Moreover, none of the ceRNA module identification methods proposed to date is able to summarize the information content of the modules in a patient- or sample-specific way. A sample-specific measure of ceRNA regulatory activity would be useful for the interpretation of its biological function and make it accessible for downstream analysis tasks such as clustering and classification. Methods that infer personalized ceRNA networks (Wang *et al*., 2022) are suited to identify individual interactions (edges) that deviate from the norm but do not capture the overall activity of ceRNAs (nodes) in the network.

Here, we introduce spongEffects, a new method to (a) extract ceRNA modules from previously inferred ceRNA networks (e.g., with SPONGE (List *et al*., 2019)) and (b) to assess their regulatory activity using enrichment scores as surrogates for module activity. These spongEffects scores (enrichment scores of the modules) are calculated at the single patient- or sample-level, thus allowing the study of ceRNA effects across groups of samples or patients or even in personalized analyses.

Given gene expression data and a pre-computed ceRNA network, spongEffects performs several steps (Fig. 1): (a) it filters for significant interactions with meaningful effect size and identifies the most important ceRNAs via network centrality analysis; (b) for the subset of nodes with the highest centrality values, it constructs ceRNA modules by incorporating first-degree ceRNA network neighbors; (c) it performs single-sample gene set enrichment using the identified nodes, thus obtaining spongEffects scores; and (d) it uses the spongEffects scores to perform downstream machine learning tasks for classification and biomarker identification.

**Figure 1:**
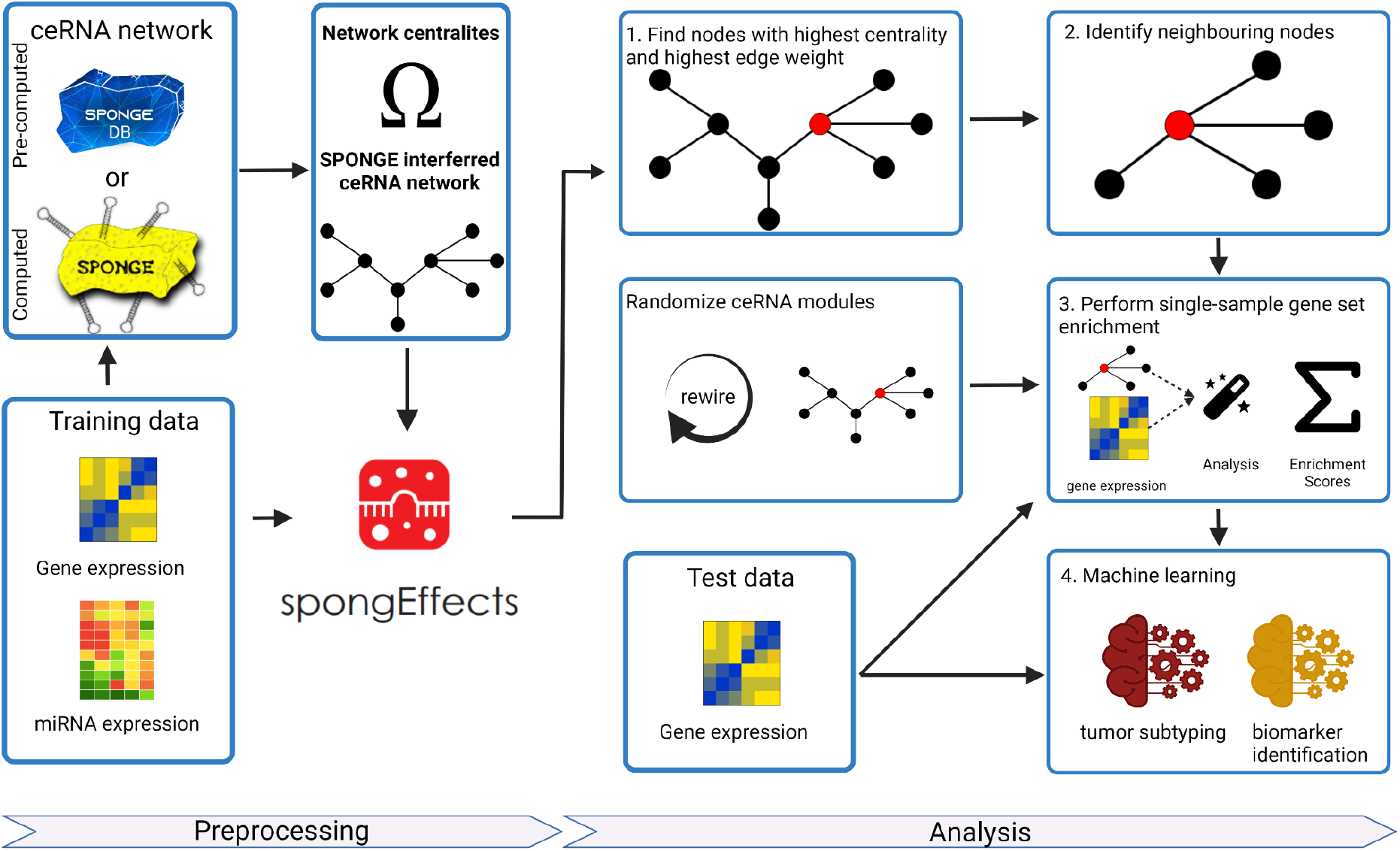
The workflow of spongEffects. spongEffects accepts a gene expression matrix and a ceRNA network as input. Subsequently, it a) filters the network and calculates weighted centrality scores to identify important nodes, b) identifies first neighbors, c) runs single-sample gene set enrichment and d) prepares the output for further downstream tasks (e.g., machine learning-based classification and extraction of mechanistic biomarkers).

We applied spongEffects to ceRNA networks inferred with two partial correlation-based methods (SPONGE (List *et al*., 2019) and Positive correlation (J. Zhang *et al*., 2019)) and one conditional mutual information-based method (Hermes (Sumazin *et al*., 2011)) and investigated how robust these modules are across different datasets. We used data from two independent breast cancer cohorts to test the enrichment methods underlying spongEffects for robustness to missing values and across different technologies (e.g., RNA-seq and microarray). We further investigated how consistent the ceRNA models are across ceRNA networks or when swapping train (i.e., the cohort used for ceRNA network inference and training a classifier) with the test data (the second independent cohort). We showed that spongEffects scores could be used to classify breast cancer subtypes with good accuracy. Finally, we show that non-coding RNAs outperform coding RNAs in classification. Importantly, once a ceRNA network is inferred, spongEffect scores can be computed from gene expression data alone, i.e., in the absence of miRNA expression profiles which are rarely available. By offering a systems biology view on the miRNA regulatory landscape, spongEffect scores are suited to uncover important ceRNAs and miRNAs in cancer biology and generate new hypotheses on the role of long non-coding RNAs.

## MATERIALS AND METHODS

spongEffects was implemented in R (version 3.6.2) and is provided to the community as a new function in the SPONGE package in Bioconductor (see Availability). As datasets, we downloaded log2(TPM + 0.001) transformed TCGA-BRCA expression data (RNAseq) from the XENA Browser (Goldman *et al*., 2020). In addition, we downloaded gene expression data (Illumina HT 12, EGAD00010000434) and miRNA expression data (Agilent ncRNA 60k, EGAD00010000438) from the 1st and 2nd METABRIC (Molecular Taxonomy of Breast Cancer International Consortium) cohorts (Curtis *et al*., 2012) from the European Genome-phenome Archive (EGA European Genome-Phenome Archive). Importantly, while the TCGA datasets contain expression values for both coding and non-coding RNAs, METABRIC contains only mRNA abundances. We used three different ceRNA inference tools to benchmark spongEffects, Positive Correlation (PC) (Xu *et al*., 2015), Hermes (Sumazin *et al*., 2011), and SPONGE (List *et al*., 2019). Given the heavy computational cost of these methods, they were run on reduced versions of the TCGA and METABRIC datasets (i.e., the 4000 mRNAs and 1000 miRNAs shared across both cohorts) with standard parameters. PC and Hermes networks were inferred with the miRspongeR Bioconductor package (J. Zhang *et al*., 2019) with standard settings. The SPONGE Bioconductor package (List *et al*., 2019) was used to calculate the SPONGE network. The same list of the candidate, ceRNA-miRNA interactions, was used to infer edges across the three methods. In order to run a more extensive analysis and analyze the role of lncRNAs in breast cancer, we further ran spongEffects on pre-computed ceRNA networks with genome-wide coverage downloaded from SPONGEdb (Hoffmann *et al*., 2021) as inputs for spongEffects. For the latter, we could also consider non-coding RNAs as ceRNA candidates which were absent in the overlap of METABRIC and TCGA.

### Defining ceRNA modules

Classical centrality measures, such as degree, closeness, and betweenness, can extract important information in biological networks by identifying hub and bottleneck nodes (He and Zhang, 2006). The definitions of such centrality measures have been generalized for applications in weighted networks (Barrat *et al*., 2004) to acknowledge that not all interactions are equally important. Here we use a weighted centrality measure composed of the following centralities:

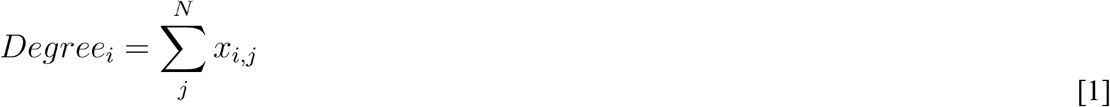

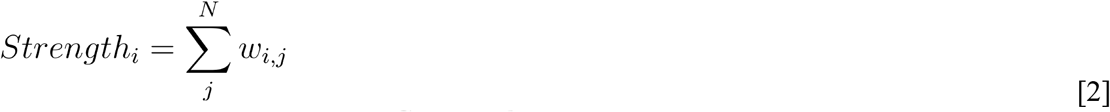

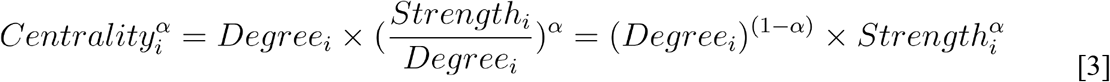

Equation [1] describes the definition of the degree of centrality of a node *i* in an undirected network of size *N*. *x_ij_* is the binary adjacency matrix describing the connection of nodes *i* and *j* (i.e., *x_ij_* = 1 if *i* and *j* are connected in the graph and 0 otherwise). Equation [2] formalizes the weighted degree centrality measure or node strength of node *i* in a network of size N. In particular, *w_ij_* > 0 if nodes *i* and *j* are connected and it corresponds to the weight of the edge between *i* and *j*. Centrality is defined in Equation [3]. Opsahl et al. describes *α* as “a positive tuning parameter that can be set according to the research setting and data. If *α* is 0 between 1 and, then having a high degree is taken as favorable, whereas if it is set above 1, a low degree is favorable” (Opsahl, 2009). spongEffects uses weighted degree centrality as implemented in the R package tnet (version 3.0.16) (Opsahl, 2009) with the alpha parameter set to 1. In the context of a ceRNA network, we thus prioritize ceRNAs with a large sum of weights (using multiple sensitivity correlation (mscor) as effect size). The mscor extends the definition of sensitivity correlation by considering the effect of multiple miRNAs for the computation of the partial correlation (see List et al. (List *et al*., 2019)). We define ceRNA modules as subnetworks consisting of ceRNA genes with the highest weighted centrality scores (central node) and their first-degree neighbors.

### spongEffects scores

We implemented three different gene set enrichment approaches to aggregate the information associated with genes belonging to the newly defined spongEffects modules into unique enrichment scores, which we call spongEffects scores. These methods belong to the class of unsupervised and single-sample assessment tools (Hänzelmann *et al*., 2013), in that they do not require initial knowledge of existing phenotype groups, are often unavailable, and yield patient-specific scores for each module. In this work, we treat the genes in a module as a gene set. The approaches include single sample Gene Set Enrichment Analysis (ssGSEA) and Gene Set Variation analysis (GSVA) algorithms as implemented in the GSVA package (version 1.34.0) (Hänzelmann *et al*., 2013), and Overall Expression (OE) (Jerby-Arnon *et al*., 2018).

Importantly, these approaches allow the calculation of the spongEffects scores even when some of the ceRNAs (e.g., circRNAs or lncRNAs) in the modules are not present in the input expression matrix, and result in module-by-sample score matrices that can be used for downstream analyses. When comparing results between GSVA, ssGSEA, and OE (Suppl. Fig. 1) we observed no major differences in the results. The original GSVA publication highlights that the choice of the optimal single-sample enrichment tool is highly dependent on the specific task and dataset and cannot be defined a priori.

spongEffects scores were calculated for the two datasets (i.e., TCGA and METABRIC) separately. Comparison of the different enrichment methods showed that scores calculated with OE, GSVA, or ssGSEA resulted in models with similar subtype classification performances (Suppl. Fig. 1). For all the subsequent analyses, spongEffects scores were calculated via OE, given its previous application in similar bulk transcriptional profiles (Jerby-Arnon *et al*., 2018).

### Machine learning for subtype classification

We trained three classes of machine learning models, namely Random Forest, Support Vector Machine with the linear kernel (Linear SVM), and eXtreme Gradient Boosting (XGBoost), as implemented in the caret R package (version 6.0.90) (Kuhn, 2008) to classify tumor samples in the respective subtypes, using spongEffects scores calculated on the training dataset as input. For each approach, we optimized the following hyperparameters via repeated (3x) 10-fold cross-validation: i) a number of features randomly sampled at each split (*mtry*) for the Random Forest models, ii) the cost (*C*) of misclassification for linear SVM models, and iii) the number of trees (*nrounds*), maximum tree depth (*max_depth*), and learning rate (*eta*) for XGBoost. We implemented a stratified approach in order to preserve the proportion of samples in each subtype at each iteration of the cross-validation procedure. Given the multi-classification problem at hand, we defined the optimal parameter as the one resulting in the highest subset accuracy (Ghamrawi and McCallum, 2005) during cross-validation. After cross-validation, we used the learned parameter to train a new classifier on the complete training set. To test these models, we predicted subtype labels using spongEffects scores calculated on a second independent expression dataset. This means that the training data set was used to infer a ceRNA network, identify ceRNA modules, and train a classifier, while the test data were not part of the training procedure and were thus considered a valid external testing dataset. The approach resulted in two performance metrics, subset accuracies for the training and testing sets, that were used to evaluate and compare the models. To aid in the interpretation and identification of modules driving subtype predictions in the Random Forest models, we calculated the Gini index as implemented in the randomForest package (version 4.6.14) (Svetnik *et al*., 2003).

### Randomization of the ceRNA networks

We designed a procedure to validate our findings and evaluate whether the modules defined by spongEffects are robust. In particular, for each original module, we sampled a random module of the same size while preserving the same size distribution of the real modules. Subsequently, the random modules are used to train a classifier as previously described. An AIMe report has been generated at https://aime-registry.org/report/UHkSyD (Matschinske *et al*., 2021).

## RESULTS AND DISCUSSION

The spongEffects method offers sample-specific insights into the miRNA regulatory landscape and highlights ceRNAs heavily engaged in miRNA cross-talk and thus implicated in many biological processes as biomarkers. To illustrate the potential of spongEffects scores for functional interpretation and machine learning, we exemplarily consider the task of breast cancer subtyping.

Breast cancer, the leading cause of cancer-related deaths in women (Sung *et al*., 2021), is a heterogeneous disease characterized by five subtypes (Luminal A (LumA), Luminal B (LumB), Basal, HER2-positive (Her2), and Normal) with different genetic prognostic profiles (Perou *et al*., 2000). Mechanisms of miRNA dysregulation play a role in breast cancer (Mulrane *et al*., 2013), offering the chance to validate miRNA mature strands as potential biomarkers (Hamam *et al*., 2017). Here, we use spongEffects to study differences in miRNA regulation at the patient-specific level based on data from The Cancer Genome Consortium (Cancer Genome Atlas Network, 2012) (TCGA-BRCA, number of samples (n) = 1,063) and the Molecular Taxonomy of Breast Cancer International Consortium (Curtis *et al*., 2012) (METABRIC cohorts 1 and 2, n = 1,905). For both cohorts, we removed all samples for which the subtype was missing or not identified as one of the five classes of interest (LumA, LumB, Her2, Basal, Normal). Furthermore, we filtered out 19 samples from the TCGA cohort for which the tumor stage indication was missing (5 patients) or unclear (Stage X, 14 patients). This step resulted in two final datasets: (1) TCGA-BRCA (n = 944), which we used for model training, and (2) METABRIC (n = 1,699), both of them were used once as training and once as an independent external validation set to build subtype classifiers as in standard machine learning applications to control for potential overfitting (see Suppl. Table 1 for the number of samples per subtype for TCGA and METABRIC).

### spongEffects scores are robust across different datasets

A wide variety of methods for the identification of potent ceRNA interactions is available in the literature (J. Zhang *et al*., 2019). These methods have been found to result in very different ceRNA networks, often sharing just a few miRNA sponge interactions (J. Zhang *et al*., 2019). We validated this result by comparing the results of three methods commonly used for ceRNA network inference: Positive Correlation (*PC*) (Xu *et al*., 2015), Hermes (Sumazin *et al*., 2011), and SPONGE (List *et al*., 2019). Given the computing time these methods take, especially for high dimensional datasets with hundreds of observations, we downsampled the TCGA and METABRIC datasets to the 4000 genes and 1000 miRNAs sequenced in both. After inference, the networks were filtered based on the statistical significance (i.e., adjusted p-value < 0.05 for Hermes and SPONGE) and strength of the inferred edges (corr > 0.1 for PC and mscor > 0.1 for SPONGE) to preserve significant associations with non-negligible effect sizes (List *et al*., 2019). This step resulted in 3773 and 1007 edges for the PC networks inferred from METABRIC and TCGA, respectively, 1103 and 60 edges for Hermes, and 3090 and 2495 edges for SPONGE. Weighted centrality measures were calculated for each remaining node in the networks (see Material and Methods). As expected, the number of interactions shared across different datasets and methods was generally quite low (Fig. 2), confirming the high dataset- and method-specificity of ceRNA network inference tools and emphasizing that further work is needed to understand the discrepancies between existing methods.

**Figure 2:**
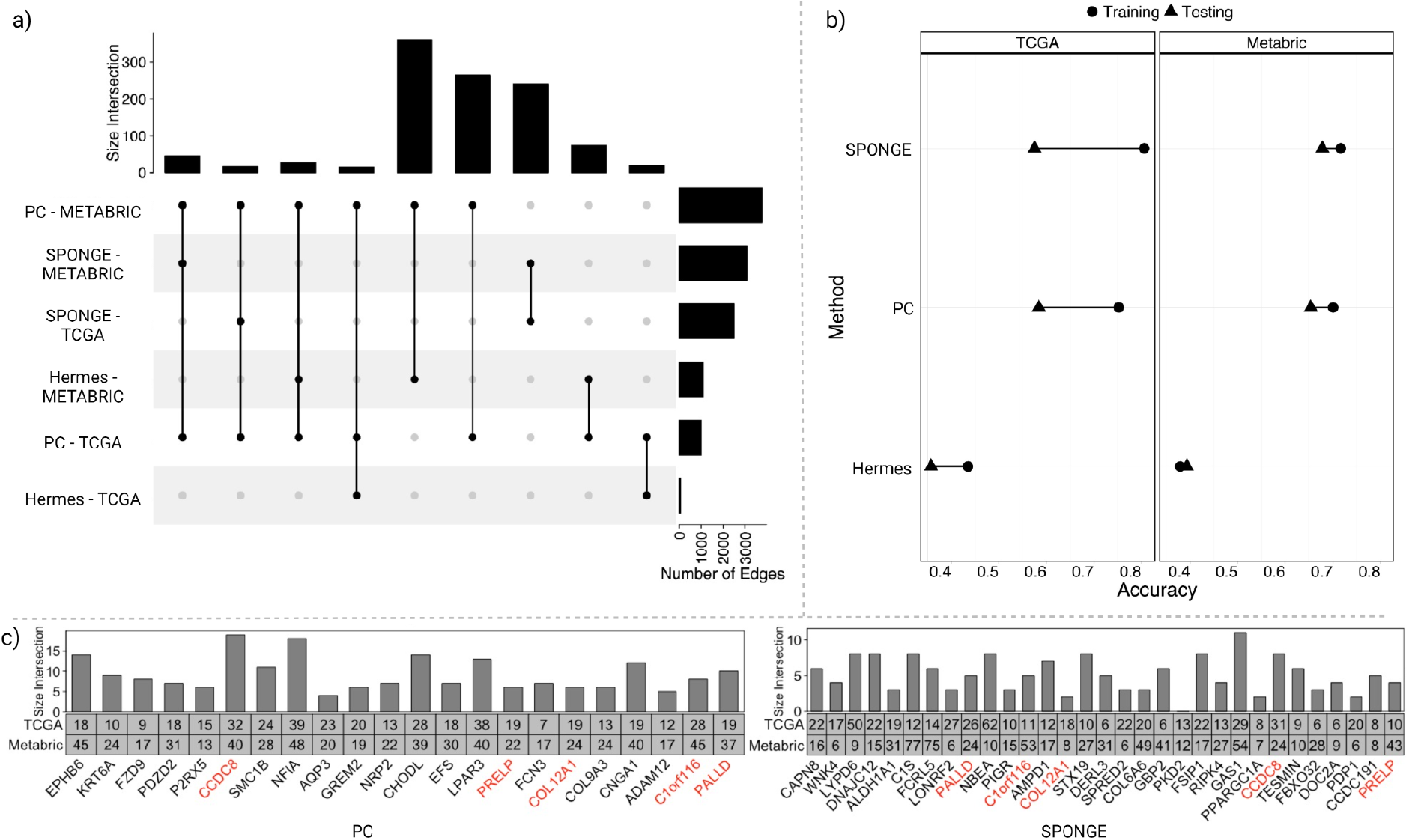
Testing the robustness of spongEffects modules using the ceRNA network inference methods PC, Hermes, and SPONGE. (a) shows the overlap of the same edges between the datasets METABRIC and TCGA-BRCA. (b) shows the accuracy of PC, Hermes, and SPONGE using either METABRIC or TCGA-BRCA as training or independent test set (e.g., if METABRIC is used as a training set, then TCGA-BRCA is used as a test set and vice versa). (c) We compared the top-scoring spongEffects modules (Gini index) from PC and SPONGE. The numbers in the boxes indicate the module size, while the bars indicate the overlap of modules obtained from training on the two different data sets. Red means that the same central ceRNA was found in both methods. We excluded Hermes from this evaluation as it achieved very low prediction performance.

To compare the predictive power of modules calculated across different datasets and methods, we trained different types of classifier models to predict breast cancer subtypes based on spongEffects scores. Random Forest, Support Vector Machine with the linear kernel (linear SVM), and eXtreme Gradient Boosting (XGBoost) models were used to build two separate families of models in order to evaluate the robustness of the modules. The first one was trained with modules calculated on networks inferred from the TGCA-BRCA dataset and validated on the METABRIC modules. In the second one, the training and datasets were switched. Upon training, we investigated the prediction performances of the model families and the similarity of the modules driving predictions in both contexts.

Despite the limited number of edges and dataset sizes, the resulting modules showed relatively good predictive performances independently of the type of machine learning model (Supp. Fig. 2). While the machine learning models built on the TCGA modules displayed higher predictive performance during training, classifiers trained on METABRIC showed better generalization capabilities and resulted in higher accuracy on the TCGA test set (Fig. 2 b). The performances of models built on modules extracted from different ceRNA network tools were quite heterogeneous. In this particular setting, the Hermes method seemed to drastically underperform when compared to modules built on PC or SPONGE modules and was thus left out for subtype prediction.

We next investigated if similar ceRNA modules were selected as top-performing classification features when training on TCGA or METABRIC data, respectively, as this would emphasize the robustness of our approach as well as the capacity of spongEffect scores to reflect important aspects of tumor biology. We ranked modules by their feature importance (mean decrease in Gini index) and investigated the overlap of the top 20 modules reported by the PC- and SPONGE-based models between the TCGA and the METABRIC-trained classifiers. 18/20 modules were identical in the PC-based models and 10/20 in the SPONGE models, indicating that the PC method is more robust in feature selection. We found only five of the top 20 modules to overlap from PC and SPONGE, which once more emphasizes that ceRNA inference methods following different concepts show comparably little agreement in results (J. Zhang *et al*., 2019).

### spongEffects modules predict breast cancer subtypes

Long non-coding RNAs have often been discussed for their relevance in breast cancer (Su *et al*., 2014) and for their role as miRNA sponges and potential biomarkers (Wang *et al*., 2022). We investigated the capacity of long-non-coding RNAs as potential ceRNAs by applying spongEffects to the TCGA-BRCA ceRNA network in SPONGEdb (Hoffmann *et al*., 2021). The original network, containing ~3 × 10^7^ edges between multiple RNA classes (e.g., mRNAs and lncRNA), was filtered down to 702,026 edges as described above. The 750 lncRNAs with the highest weighted centrality measures were selected and used as central nodes to define the spongEffects modules. We retain modules containing 10 to 200 genes after removing genes missing in the expression matrices, as previously suggested (Hänzelmann *et al*., 2013).

Fig. 3 shows that the distribution of spongEffects scores overall found modules differ markedly between subtypes. We note that the distribution of the basal subtypes diverges from a normal-like distribution suggesting a hidden subgroup of patients. After dissecting this distribution using the R package mixtools (version 1.2.0) (Benaglia *et al*., 2009), we find that the smaller group is enriched in stroma-high tumors (Fig. 3 and Suppl. Fig. 3.a), as calculated via ESTIMATE (Yoshihara *et al*., 2013), and in extracellular matrix (ECM)-related genes, which have been associated with Basal invasion programs under the regulation of KRT14 related genes (Hanley *et al*., 2020) (Suppl. Fig. 3.b). Analogous studies have identified similar Basal subgroups from different data types and cohorts (Asleh *et al*., 2022). This highlights the ability of spongEffects to identify a subset of samples with potential prognostic values.

**Figure 3:**
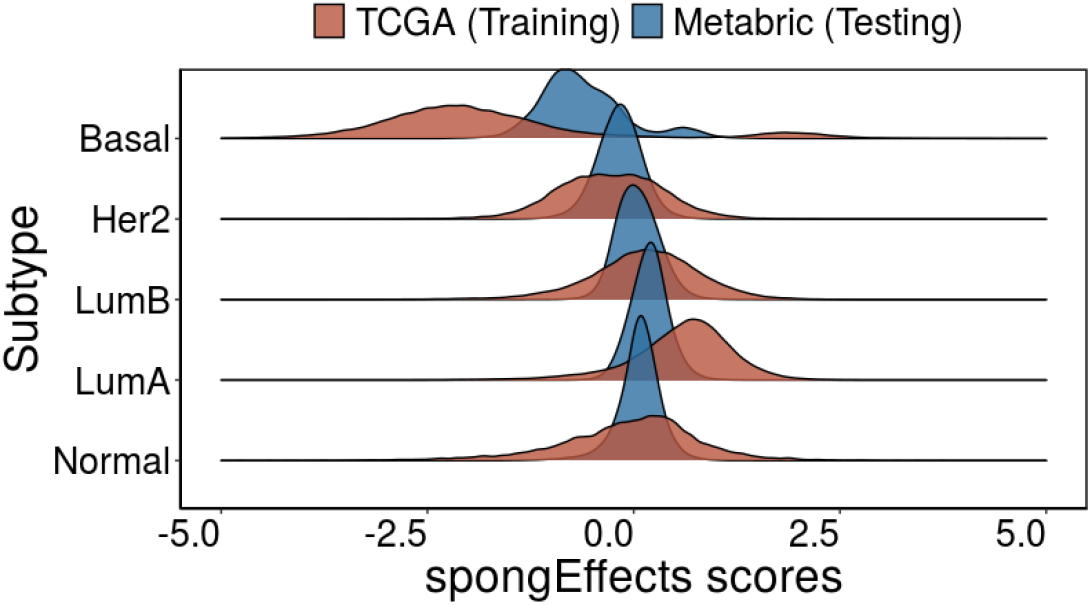
Distribution of the spongEffects scores over all modules in the TCGA-BRCA (training) and METABRIC (testing) dataset. With the exception of Basal breast cancer, the scores for the various breast cancer subtypes are approximately normally distributed. For the Basal subtype, the bimodality of the distribution hints at the existence of a subgroup of patients with characteristics different from the rest of the class (Suppl. Fig. 3).

We trained a Random Forest classifier on the TCGA-BRCA dataset and validated the resulting model on the METABRIC dataset. We repeated this analysis using different numbers of lncRNAs (200, 250, 500, 750, 1000, 1500, and 2000) to study the robustness of our approach and to highlight the relatively small influence of the number of initial lncRNAs (and thus modules) considered on the final results (Suppl. Fig. 4). To assess whether spongEffects scores offer additional value over the expression level of the ceRNAs alone, we compared our results to the predictive performance of randomly defined modules and a baseline model trained on the expression values (not on spongEffects scores) of the 25 lncRNAs common to both datasets. The subset accuracy metric (Ghamrawi and McCallum, 2005) was used to evaluate the classification performance along with sensitivity, specificity, and F1 score (confusion matrices in Suppl. Fig. 5). The spongEffects-based classification model outperformed the random module-based and the lncRNA-based models both in training and testing (Fig. 4.a) and preserved good specificity, sensitivity, and F1 scores across all breast cancer subtypes (Suppl. Fig. 6.b). Moreover, a comparison of the prediction performances of models based on mRNA expression only (Fig. 2) and lncRNA-based (Fig. 4) models suggests that non-coding RNAs might carry important information about the post-transcriptional regulatory wiring in different breast cancer subtypes. Improved performance might be due to the increased number of interactions inferred when different classes of RNA are included, offering a better description of the regulatory landscape of the disease in analysis, as hypothesized in (Gysi and Barabasi, 2022).

**Figure 4:**
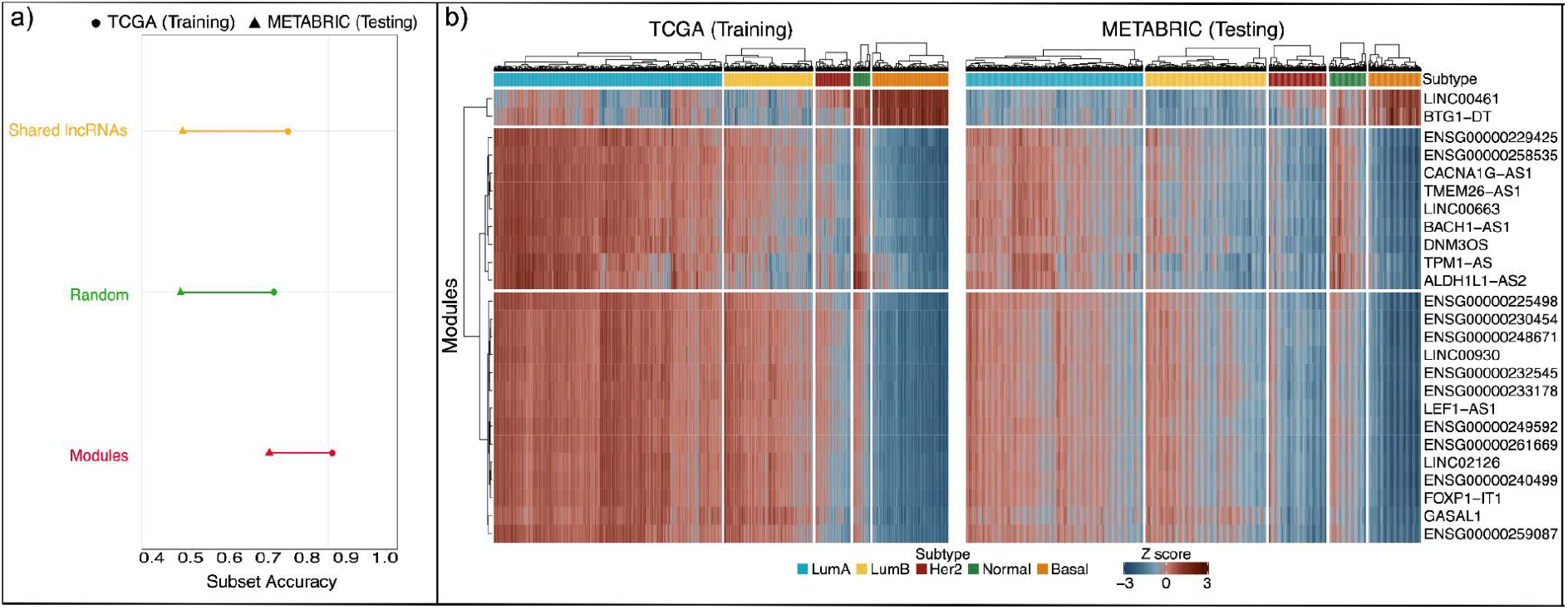
Overview of Results. Overview of performance metrics for the Random Forest models calibrated on spongEffects modules (red), randomly defined modules (green), and lncRNAs common to both the TCGA (training) and METABRIC (testing) datasets (yellow). Subset accuracy (the proportion of samples that have all their labels classified correctly) values in training and testing for the three models. b) Heatmaps showing the standardized spongEffects scores for the top 25 most predictive modules calculated on the TCGA and METABRIC datasets. The heatmaps show a clear separation of the Basal subtype from the remaining ones, potentially pointing to the role of miRNA-regulated modules in the aggressive and proliferative phenotype observed in this subtype.

### ceRNA modules identify key biological mechanisms

Upon closer inspection of the top 25 ceRNA modules across breast cancer subtypes in both datasets (Fig. 4. b, Suppl. Fig. 7), we observed that Basal samples showed lower scores for modules centered around lncRNAs that were previously suggested to act as miRNA sponges. For example, CACNA1G-AS1 was shown to decrease proliferation, EMT transition, and migration in hepatocellular carcinoma (Yang *et al*., 2019) and non-small cell lung cancer (Yu *et al*., 2018) and to downregulate p53 levels in colorectal cancer (Wei *et al*., 2020). In contrast, DNM3OS has been shown to promote tumor progression and SNAIL-mediated EMT in gastric cancers (Wang *et al*., 2019) and to confer resistance to radiotherapy by mediating DNA damage response mechanisms in esophageal squamous cell carcinoma. TPM1-AS has been shown to regulate alternative splicing of TPM1 and regulate TPM1-mediated migration of cancer cells in esophageal cancer (Huang *et al*., 2017). Modules such as LINC00461 show strong enrichment in Basal samples. LINC00461 has been shown to promote migration and invasion in breast cancer (H. Zhang *et al*., 2019; Dong *et al*., 2019) and to play a role in the progression of other cancer types, such as hepatocellular carcinoma (Ji *et al*., 2019) and renal cell carcinoma (Chen *et al*., 2019), or in mediating radiosensitivity in lung adenocarcinoma (Hou *et al*., 2020). However, such clear modules are only available for the Basal subtype, and we cannot identify such modules for the rest of the breast cancer subtypes as we believe that these are combinatorial effects of more than one module that separates the rest of the subtypes (Fig. 4. b). In TCGA-BRCA, where patient-matched miRNA and gene expression data are available, we ranked miRNAs based on how many times they were predicted by SPONGE to target the genes in the modules and noted several miRNA families shared across modules (Suppl. Fig. 8.a). 13 of the 51 miRNAs that were frequently associated with modules were also considered highly predictive in a subtype classification model based on miRNA expression data from the TCGA-BRCA dataset (Suppl. Fig. 9) and showed clear differences in expression between subtypes (Suppl. Fig. 8.b). This highlights how our approach offers insights into miRNA and ceRNA regulation even when miRNA expression data is unavailable. Focusing on the miRNA expression differences between subtypes offers a potential explanation for the spongEffects scores we observed.

### Predictive spongEffects modules can identify potential biomarkers in breast cancer

Here, as an example, we describe in more detail the CACNA1G-AS1 module and its miRNAs, which have been experimentally validated. Specifically, we investigate its role in the Basal subtype, one of the most aggressive breast cancer subtypes for which effective targeted therapies are still missing (Foulkes *et al*., 2010; Lehmann *et al*., 2011).

Genes in module CACNA1G-AS1 have lower expression levels in the Basal subtype when compared to the other subtypes in TCGA-BRCA and METABRIC (Suppl. Fig. 8.a and Suppl. Fig. 8.b) and contribute to the biology of this subtype. For example, expression of TBC1D9 has been shown to be inversely correlated to proliferation and grading in Basal breast cancer (Kothari *et al*., 2021), while genes such as MYB and ZBTB16 are tumor suppressors whose promoter regions are often hypermethylated in Basal breast cancers (He *et al*., 2020; Roll *et al*., 2013).

In order to check whether we could exploit our modules to explain such changes, we further investigated the most frequent miRNAs in the CACNA1G-AS1 module (Suppl. Fig. 8) and checked their expression in the TCGA-BRCA dataset. Interestingly, miR-301b-3p and miR-130b-3p had higher expression in the Basal subtypes, giving a potential explanation for the spongEffects score of the genes they target (Suppl. Fig. 10.c). The same approach can be applied to module LINC00461, where genes belonging to these modules are known to be upregulated in Basal cancers and contribute to its highly proliferative and aggressive phenotype (Suppl. Fig. 10.b and Suppl. Fig. 10.d).

Additionally, we investigated if the members of a module jointly contribute to important biological functions via gene set enrichment analyses with g:Profiler (Raudvere *et al*., 2019). The majority of modules were enriched in terms describing functions of the extracellular matrix, plasma membrane, and cell-to-cell signaling (Suppl. Excel Tables). These results appear reasonable since it was previously shown that the state of the extracellular membrane could predict the prognosis of the patient (Robertson, 2016). Moon et al. showed in experiments with the MCF-7 cell line that there are changes in the plasma membrane between subtypes (Moon *et al*., 2020), and Worsham et al. have shown that cell-to-cell signaling differentiates between breast cancer subtypes (Worsham *et al*., 2015). We further found three modules (LINC02126, ENSG00000240499, FOXP1-IT1) that are enriched for the term mammary gland development, a crucial process in breast cancer that is known to be deregulated (Suppl. Excel Tables) (Chen *et al*., 2021; Ercan *et al*., 2011). Two other modules are enriched in known cancer pathways (DNM3OS, TPM1-AS) (Suppl. Excel Tables).

## CONCLUSION

spongEffects is a new method for extracting and interpreting ceRNA modules from pre-computed ceRNA networks. To the best of our knowledge, spongEffects is the first method that offers a sample- or patient-specific score that can quantify the regulatory activity of a ceRNA and its associated miRNAs. We demonstrated how the spongEffects method could be applied to ceRNA networks inferred with different approaches. Considering the use case of breast cancer subtype prediction, we could show that spongEffects module scores are robust features for subtype classification across independent cohorts and gene expression profiling technologies. Our analysis confirmed previous studies highlighting low agreement between ceRNA inference tools and the need for further research into method benchmarking and possibly the use of consensus strategies. While this is an active area of research, we see spongEffects as a way to identify ceRNA modules generalizable across different cohorts and suitable as potential biomarkers.

We expect spongEffects scores to reflect two overlapping regulatory mechanisms. The first is miRNA-mediated ceRNA-target regulation, where an increase in the expression of a ceRNA leads to miRNA depletion, relieving other target genes from miRNA repression and, conversely, a decrease in ceRNA expression leaves more miRNA copies that can act as repressors of their target genes (Fig. 5 middle). The second mechanism is the classical miRNA regulation, where an increase in the expression of a miRNA leads to the repression of its target genes and vice versa (Fig. 5 bottom). While it is possible to verify the former in specific settings, e.g., in the presence of miRNA expression data, it is challenging to disentangle the two effects computationally. Single-sample enrichment tools estimate the enrichment or depletion of the spongEffects modules with respect to the expression of all the other genes in the expression matrix. In spite of the difficult interpretation, spongeEffects scores highlight important biomarkers. We have calculated spongEffects scores for samples from 22 different cancer types from TCGA and made the scores publicly available so that the community can use them to infer ceRNA and miRNA regulatory activities and gain insights into the gene regulatory landscape where the cohort size is too small for robustly inferring ceRNA networks or where supporting miRNA expression data is missing.

**Figure 5:**
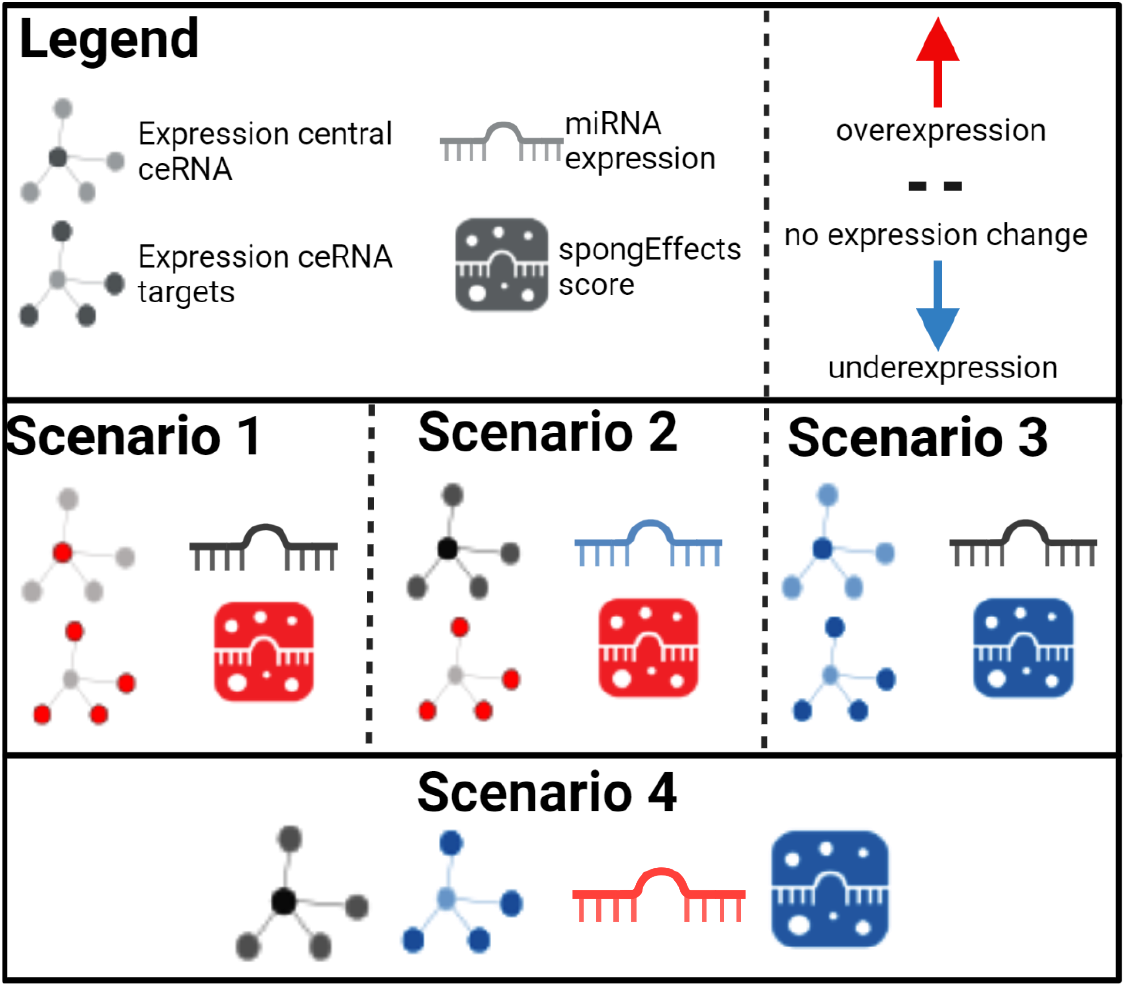
Understanding increments and reductions in spongEffects scores. Increments in spongEffects scores can result from Scenario 1) an increase in the expression of a module’s central ceRNA while miRNA expression levels remain constant, leading them to upregulation of the module’s target ceRNAs, and/or Scenario 2) a decrease in the expression of a module’s miRNAs that results in higher expression of the module ceRNAs. Negative spongEffects scores can result from Scenario 3) a decrease of expression in the module’s central ceRNA while miRNA expression levels remain constant, leading to the downregulation of the module’s target ceRNAs, and/or Scenario 4) an increase of expression in a module’s miRNAs that results in lower expression of the module’s target ceRNAs.

Using SPONGE-inferred ceRNA networks and RNA-seq data for TCGA breast cancer as inputs, we demonstrate how spongEffects scores can capture important aspects of cancer biology and be used in downstream machine learning tasks such as subtype prediction and biomarker identification. Using independent microarray-based gene expression data from the METABRIC consortium, we show that the inferred modules generalize to different cohorts. A particular strength of our approach is that spongEffects modules, once inferred from a ceRNA network, can also be computed for datasets or cohorts that lack miRNA data. For the modules that were predictive in subtype classification, spongEffects scores capture contributions of two different regulatory mechanisms: ceRNA regulation and miRNA regulation (see Section spongEffects scores). While it is challenging to separate the two, we observe that, in practice, our method returns a robust representation of microRNA-mediated regulatory effects.

We presented an extensive use case on the application of spongEffects to study lncRNA-mediated miRNA regulation. Our analysis showed that ceRNA interactions calculated on coding and non-coding RNAs together have higher discriminative power than interactions calculated on mRNAs alone, probably due to the extended interaction space allowed by the integration of multiple classes of RNAs. Many gene regulatory mechanisms of lncRNAs are poorly understood and must be experimentally discovered (Statello *et al*., 2021). spongEffects modules could help prioritize lncRNAs that have an expected impact on cancer biology and miRNA regulation, informing experimental validation and unraveling currently uncharacterized gene regulatory mechanisms of lncRNAs in combination with miRNAs. Of particular interest here is to address the question of lncRNAs’ modes of action. For instance, it is currently debated if lncRNAs are exported outside of the nucleus, which would be a requirement for the established Argonaute-dependent mechanism of miRNA regulation (McGeary *et al*., 2019). lncRNAs with an experimentally verified role as ceRNAs could then serve as important biomarkers and potential therapeutic drug targets. We showed that our method is generally applicable to different ceRNA networks, but we envision it as a potential solution to analyze gene-regulatory networks where similar concepts have already been explored (Castro *et al*., 2016; Chagas *et al*., 2019). We foresee two interesting directions for further research. First, recent studies on the inference of regulons (Castro *et al*., 2016; Fletcher *et al*., 2013) offer the chance to integrate methods for the combined analysis of transcription factors and miRNA regulation. Second, the surge of single-cell sequencing technologies allows the study of regulatory mechanisms at a higher resolution than what was possible before (Van de Sande *et al*., 2020; Ma *et al*., 2020; Gibbs *et al*., 2022), paving the way to the design of methods with the potential of uncovering new insights into the complexity of tumor biology.

## Supporting information

gProfiler results of the top 25 modules

## DATA AVAILABILITY

SPONGE and spongEffects are available via Bioconductor and GitHub:

https://bioconductor.org/packages/devel/bioc/html/SPONGE.html

https://github.com/biomedbigdata/SPONGE

SPONGEdb is available at:

http://sponge.biomedical-big-data.de/

## ACKNOWLEDGMENTS

We want to thank Christina Trummer for designing the spongEffects logo. We further thank Marcel H. Schulz for constructive feedback.

The results published and shown here are in whole or part based upon data generated by the TCGA Research Network: https://www.cancer.gov/tcga

Images were created with https://www.biorender.com. Parts of the images were designed using resources from Flaticon.com.

## SUPPLEMENTARY MATERIAL

Supplementary material is available at *Bioinformatics* online.

