## Supplementary Figures, Tables, and Materials for "spongEffects: ceRNA modules offer patient-specific insights into the miRNA regulatory landscape"


#### ABSTRACT


**Motivation:** Cancer is one of the leading causes of death worldwide. Despite significant improvements in prevention and treatment, mortality remains high for many cancer types. Hence, innovative methods that use molecular data to stratify patients and identify biomarkers are needed. Promising biomarkers can also be inferred from competing endogenous RNA (ceRNA) networks that capture the gene-miRNA gene regulatory landscape. Thus far, the role of these biomarkers could only be studied globally but not in a sample-specific manner. To mitigate this, we introduce spongEffects, a novel method that infers subnetworks (or modules) from ceRNA networks and calculates patient- or sample-specific scores related to their regulatory activity.
**Results:** We show how spongEffects can be used for downstream interpretation and machine learning tasks such as tumor classification and for identifying subtype-specific regulatory interactions. In a concrete example of breast cancer subtype classification, we prioritize modules impacting the biology of the different subtypes. In summary, spongEffects prioritizes ceRNA modules as biomarkers and offers insights into the miRNA regulatory landscape. Notably, these module scores can be inferred from gene expression data alone and can thus be applied to cohorts where miRNA expression information is lacking.
**Availability:** https://bioconductor.org/packages/devel/bioc/html/SPONGE.html
**Contact:**
**Supplementary information**: Supplementary data are available at Bioinformatics online.

#### SUPPLEMENTARY FIGURE 1

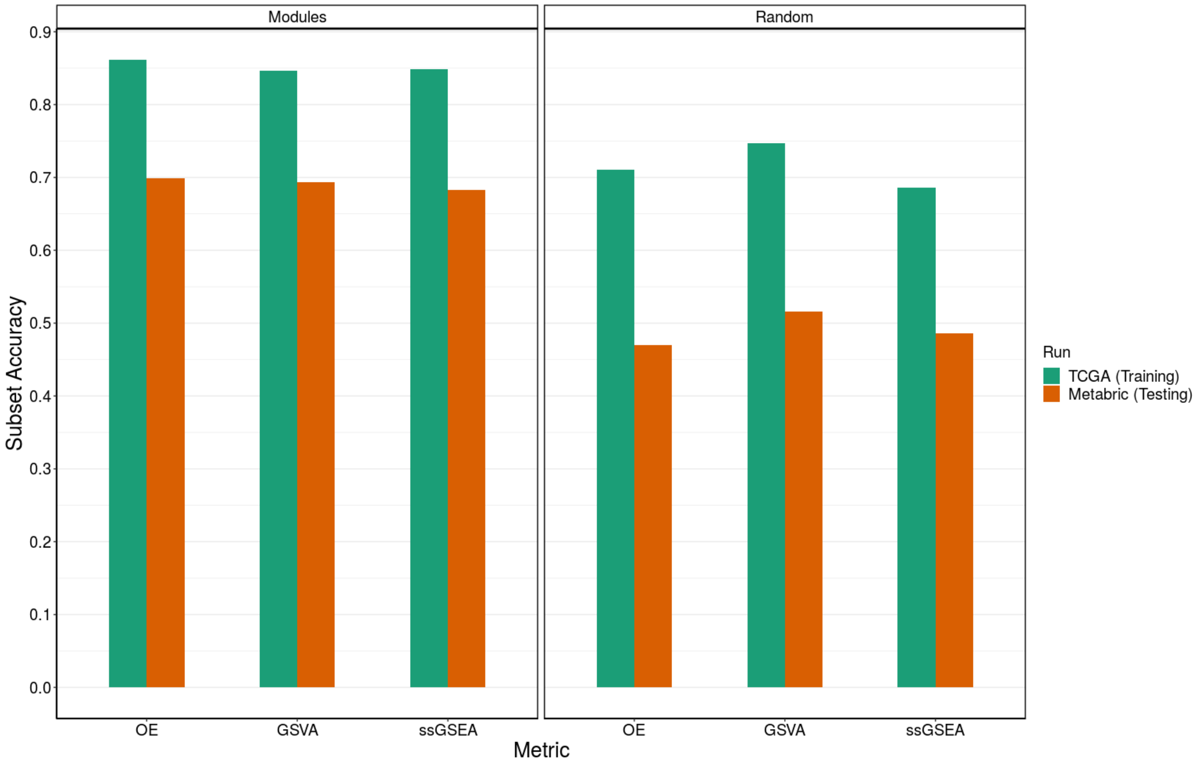


Comparison of performances in training (green) and testing (orange) of models built on spongEffects calculated using the three different single-sample enrichment tools offered by the package: Overall Expression (OE), Gene Set Variation Analysis (GSVA), and Single-Sample Gene Set Enrichment Analysis (ssGSEA). Performances were evaluated for scores calculated on spongEffects modules (left) and randomly defined groups of genes (right).

#### SUPPLEMENTARY FIGURE 2

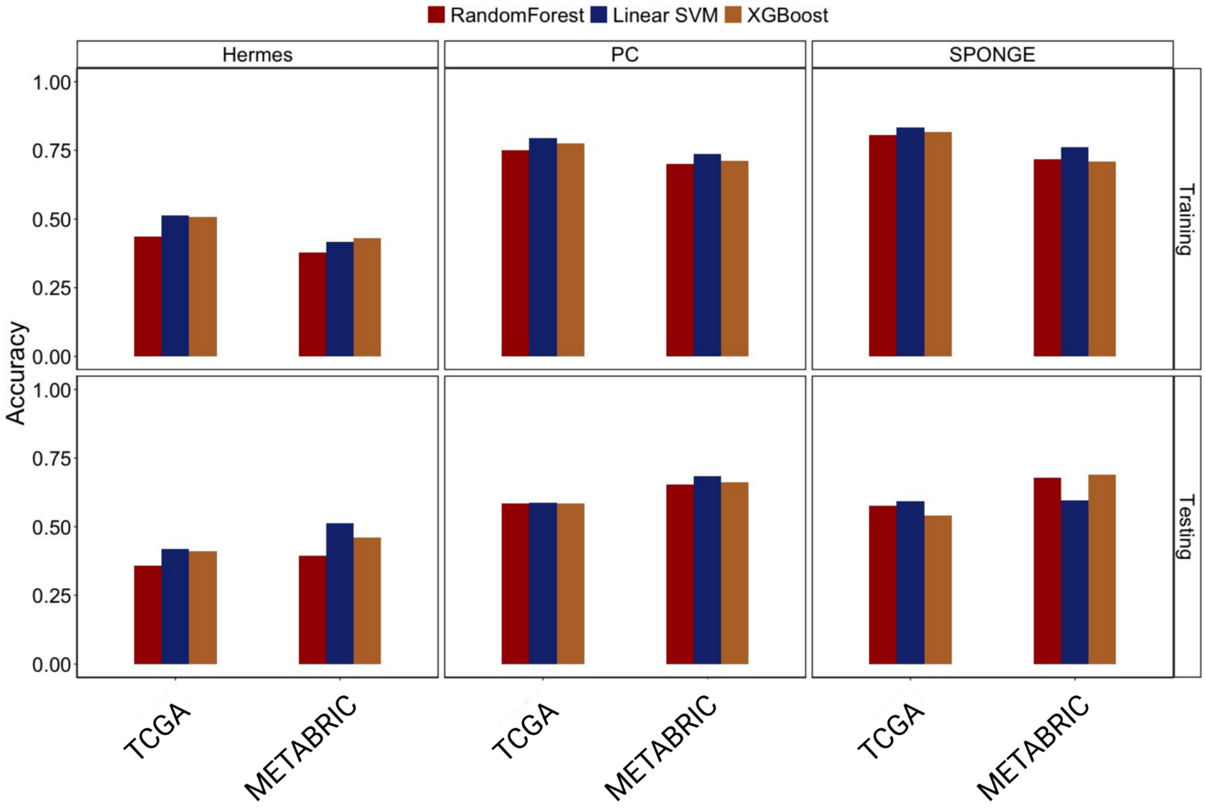


We show the accuracy of the random forest, linear SVM, and XGBoost for using TCGA and METABRIC as training and test sets.

### SUPPLEMENTARY FIGURE 3

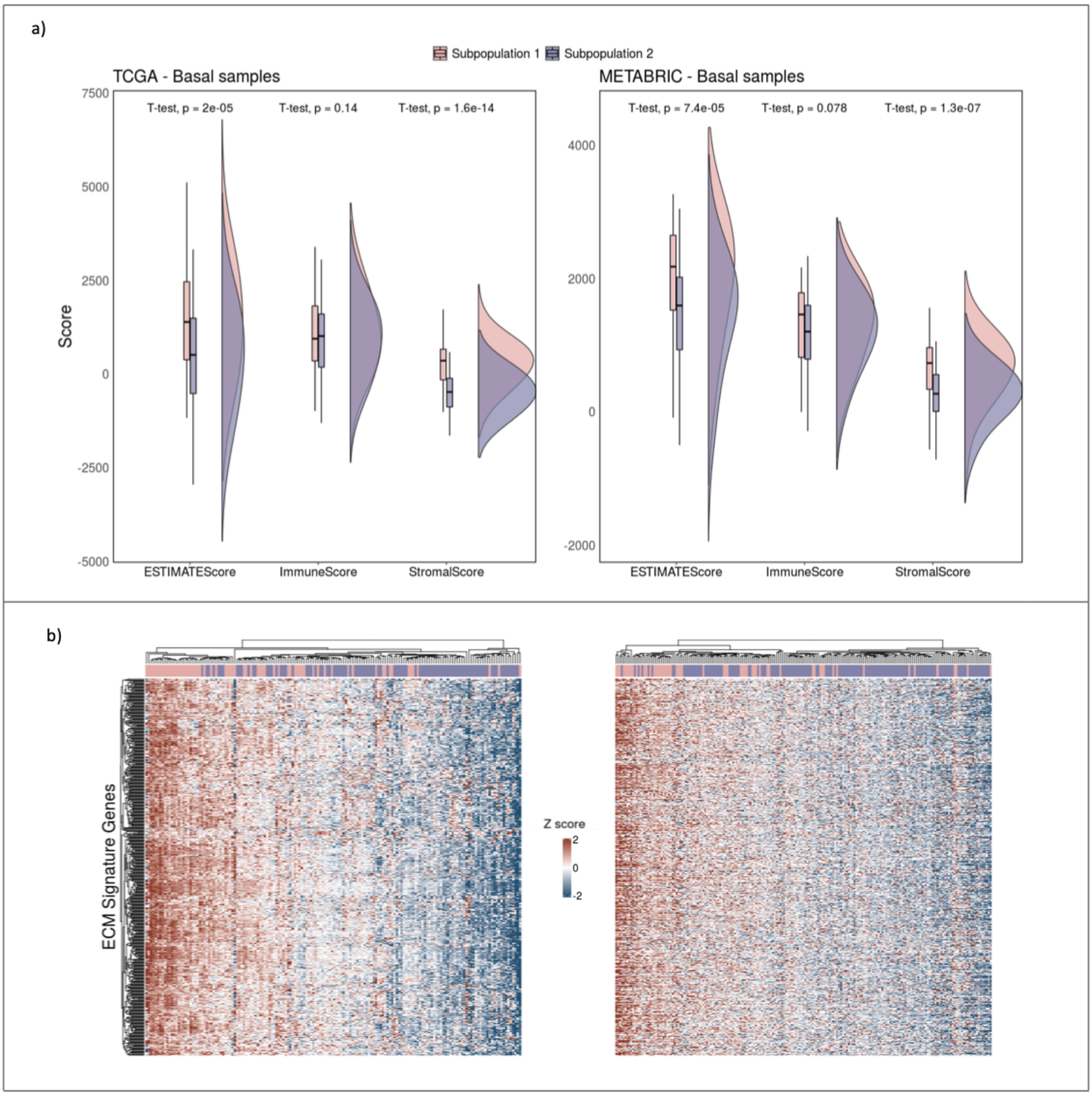


Model-based clustering applied independently to the spongEffects scores calculated for the basal samples in the TCGA BRCA and METABRIC cohorts identifies two subpopulations of patients. The differences between the two can be linked to disparities in purity, stromal content, and expression of extracellular matrix (ECM)-related genes. a) Distribution of the three types of immune scores calculated via ESTIMATE for the two identified basal populations in TCGA (left) and METABRIC (right) samples. The differences were statistically significant for scores related to purity and stromal content, highlighting potential differences in the role of miRNA regulation in the crosstalk between tumors and their microenvironment. b) Heatmaps showing the expression of ECM-related genes in TCGA (left) and METABRIC (right) basal samples. The samples previously assigned to subpopulation 1 and enriched in stroma show higher expression of the ECM signature, which is known to play a role in the development of more aggressive breast cancer phenotypes.

#### SUPPLEMENTARY FIGURE 4

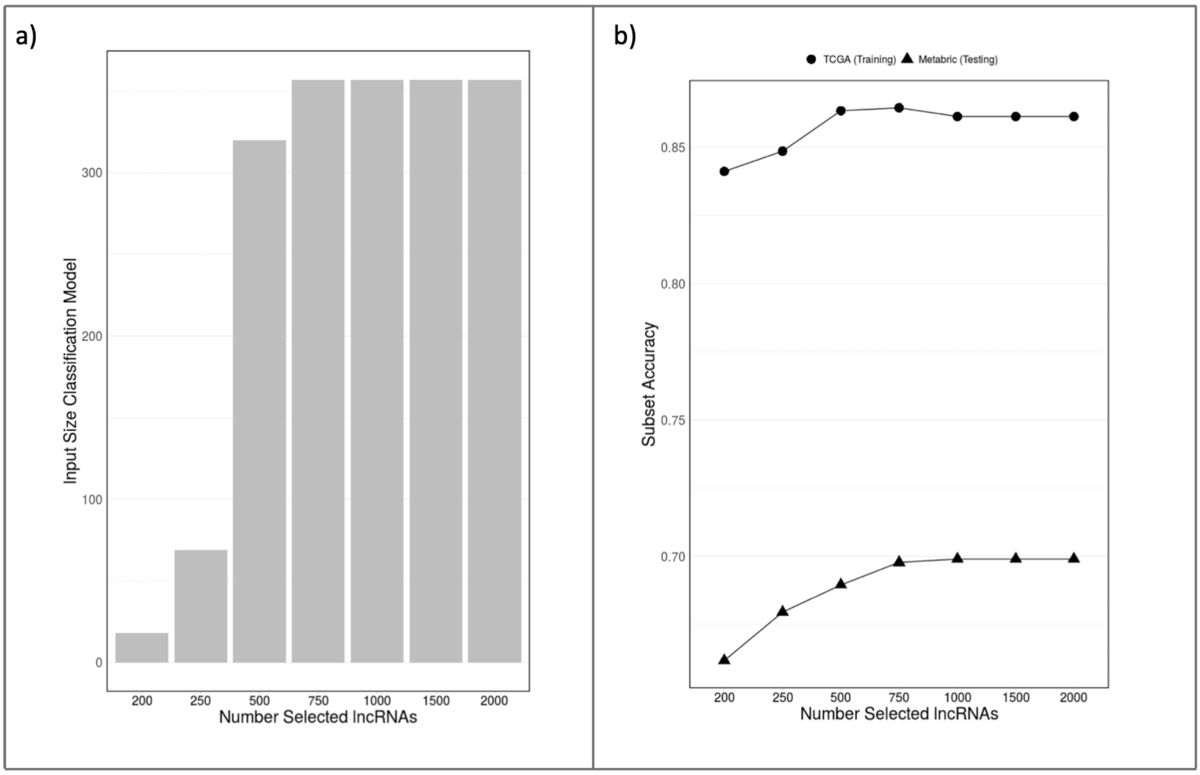


Evaluation of the optimal number of lncRNAs with high centrality to be used as an input to define the spongEffects modules and build accurate subtype classification models. a) The number of modules actually used for the classification model, after filtering for modules with more than 10 and fewer than 200 tumor samples, as a function of the initial number of selected lncRNAs. We observe that for selecting more than the top 750 lncRNAs, no additional modules pass the size filter in this data set. b) We built modules using the top 200, 250, 500,750,1000, 2000 lncRNAs with the highest weighted centrality scores and evaluated their performance on the same classification task described in the manuscript. Accuracy plateaus were reached both in training and testing when more than 750 central lncRNAs were used to define the spongEffects modules.

#### SUPPLEMENTARY FIGURE 5

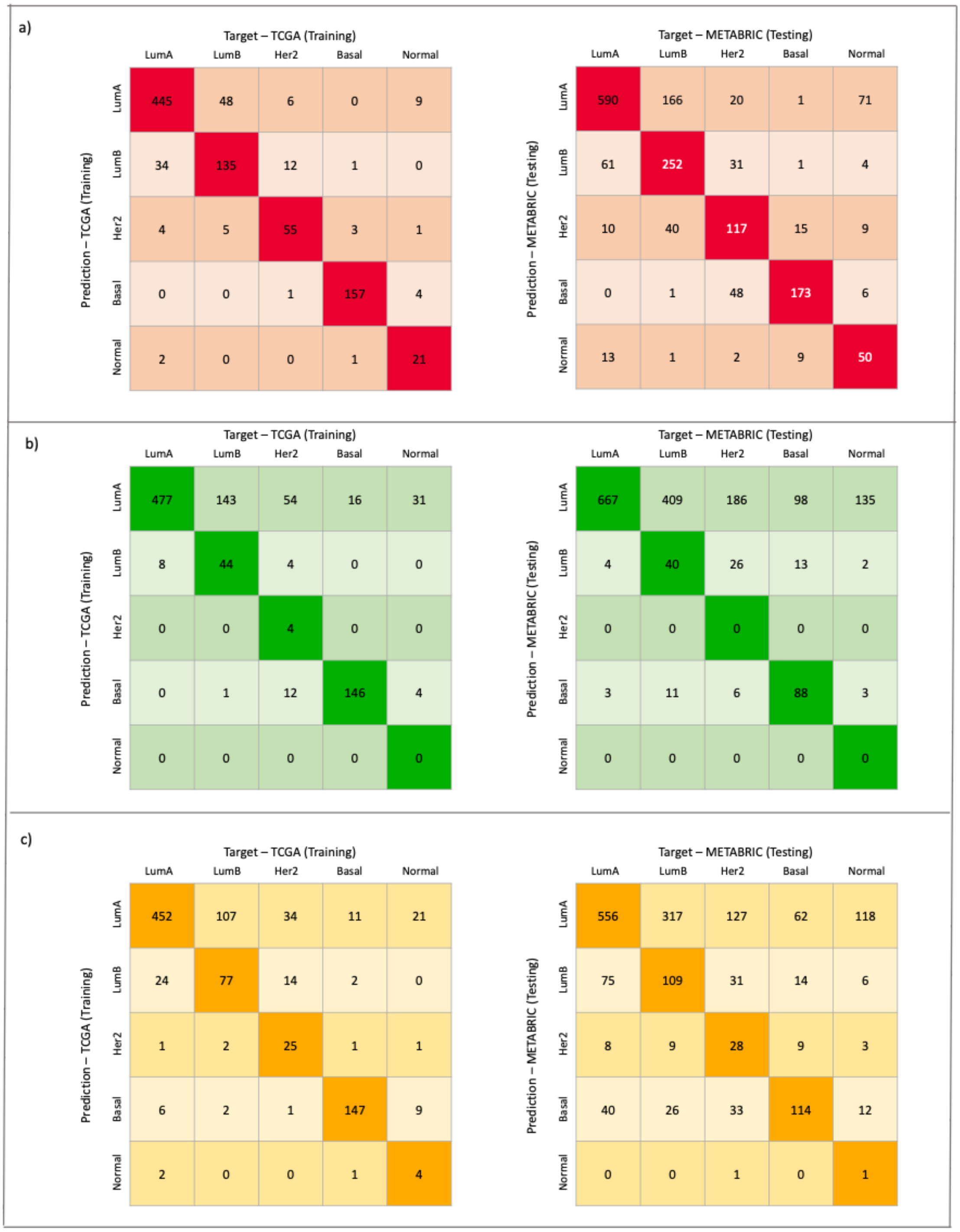


Confusion matrices representing the predictive performances of the Random Forest models on the training (TCGA, left) and testing (METABRIC, right) datasets when different inputs (from the top, spongEffects scores, random module scores, and central genes) are used. a) Confusion matrices showing the results of the classification model trained on spongEffects scores calculated for the spongEffects modules. b) Confusion matrices showing the results of the classification model trained on spongEffects scores calculated for randomly defined modules. c) Confusion matrices showing the results of the classification model trained on the expression of lncRNAs measured in both the TCGA and METABRIC cohorts.

#### SUPPLEMENTARY FIGURE 6

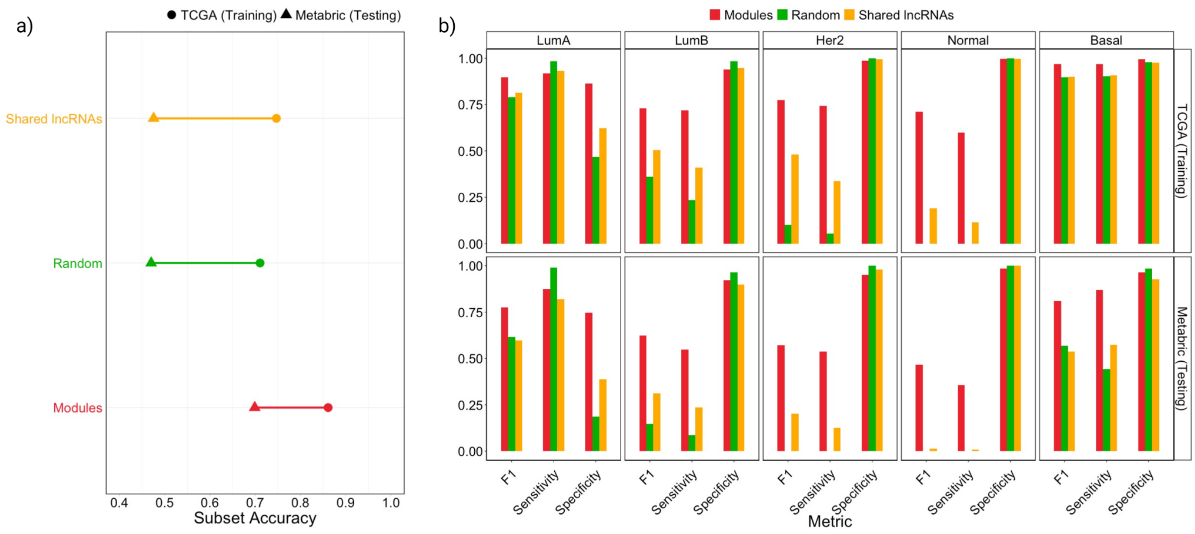


Overview of performance metrics for the Random Forest models calibrated on spongEffects modules (red), randomly defined modules (green), and lncRNAs common to both the TCGA (training) and METABRIC (testing) datasets (yellow). a) Subset accuracy (the proportion of samples that have all their labels classified correctly) values in training and testing for the three models. b) Sensitivity, specificity, and the harmonic mean of both (F1) for the three models across breast cancer subtypes.

#### SUPPLEMENTARY FIGURE 7

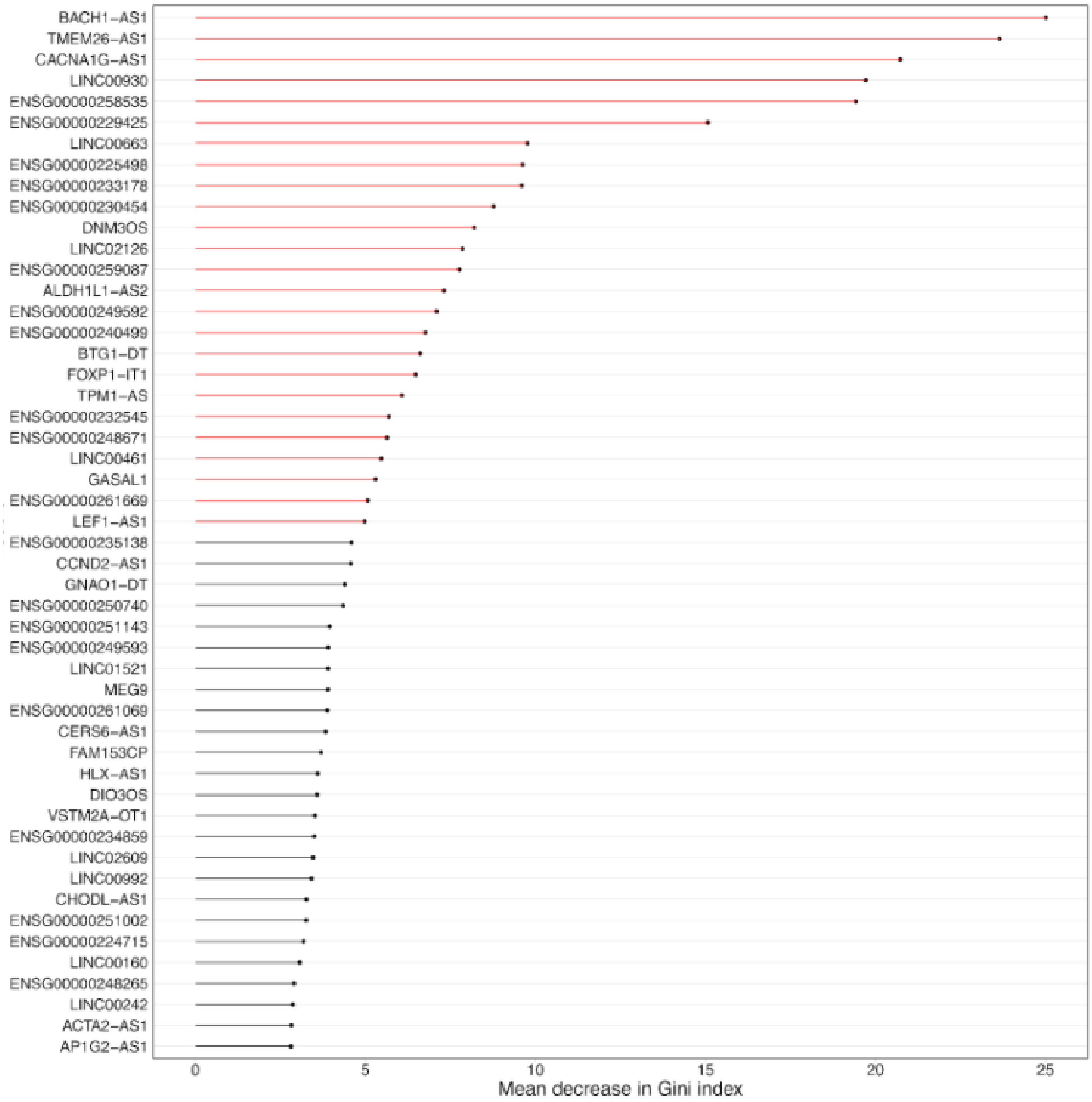


Most-predictive modules ranked by the Gini index. We selected the top 25 most predictive modules (red) for downstream analysis.

#### SUPPLEMENTARY FIGURE 8

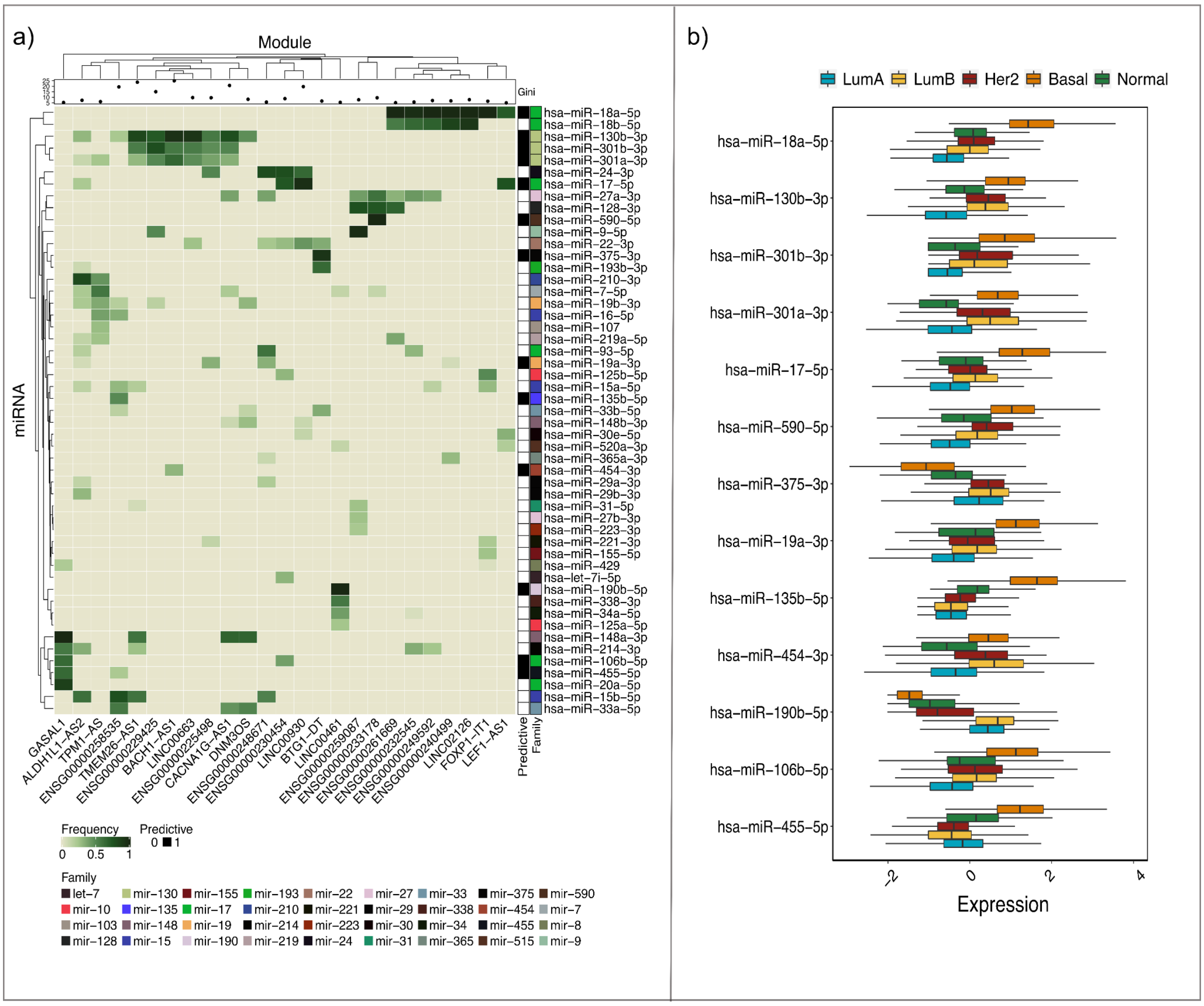


spongEffects scores can be partially explained by miRNA regulation. a) Fraction of a module’s genes that are targeted by the miRNA in the top 25 ceRNA modules. Additionally, miRNA families are shown to indicate miRNAs with a shared seed sequence. We further indicate which of the miRNAs are predictive of breast cancer subtypes. b) Expression levels of the miRNA mature strands driving classification of breast cancer subtypes.

#### SUPPLEMENTARY FIGURE 9

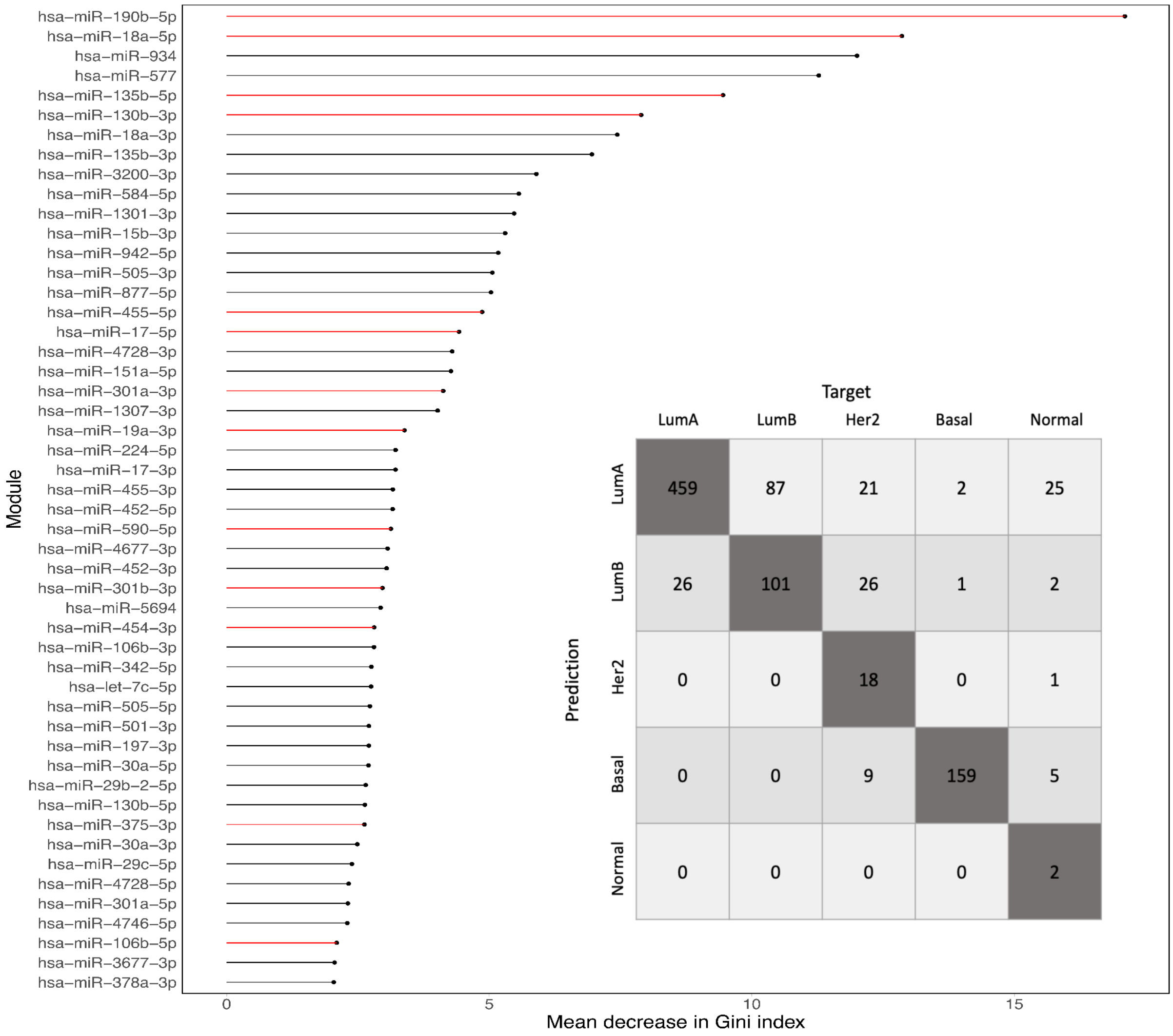


Results of the classification model trained on miRNA expression from the TCGA dataset. The most-predictive miRNAs are ranked by the Gini index. The miRNAs highlighted in red are ones predicted to regulate the genes part of the important modules described in the main text. In gray, the confusion matrix related to the performance of the miRNA-based classification model on the training (TCGA) cohort.

### SUPPLEMENTARY FIGURE 10

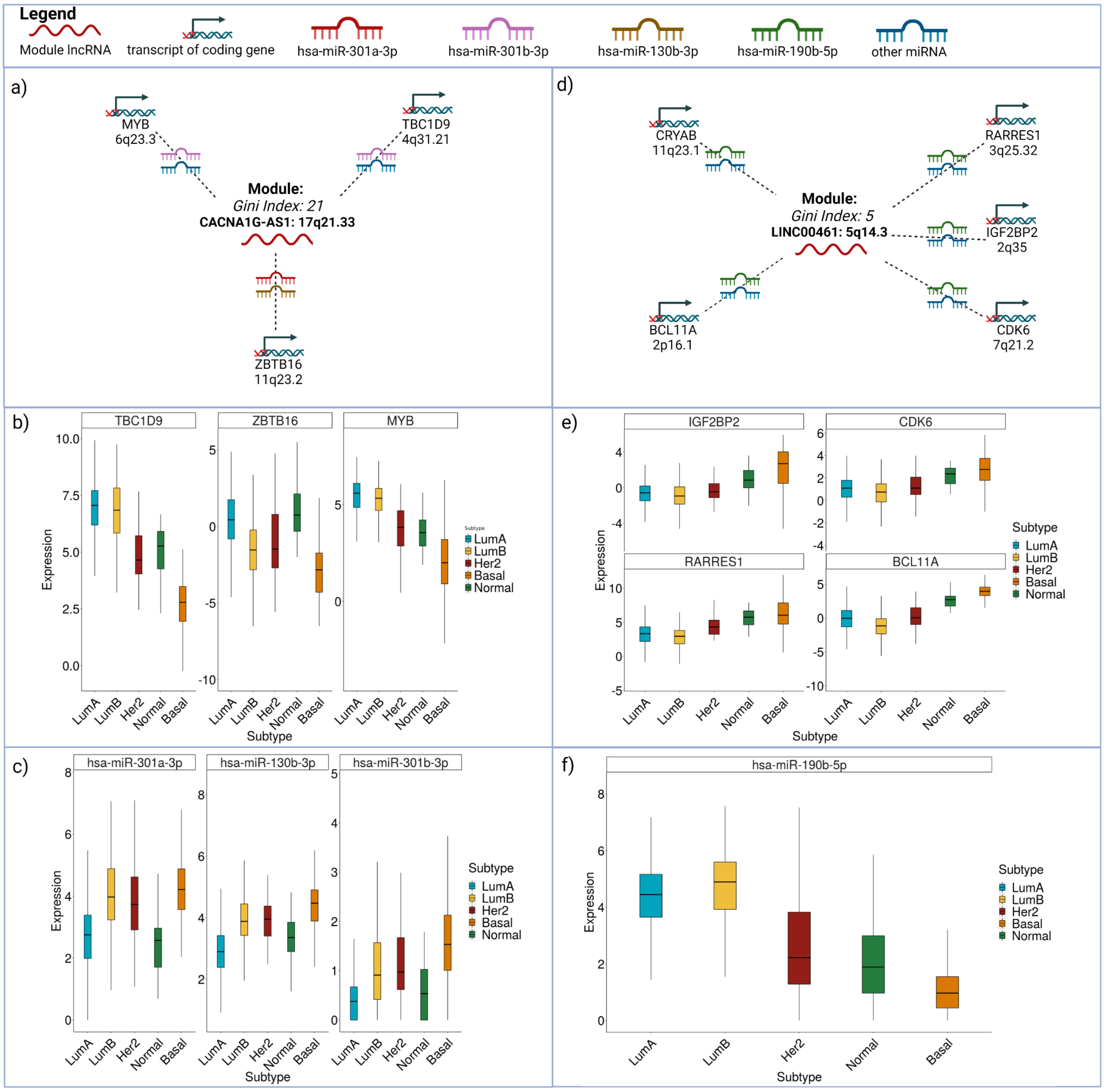


Expression of genes in the CACNA1G-AS1 and LINC00461 modules and miRNAs predicted to target them. These genes have been experimentally validated to play a role in Basal breast cancers. a) Three genes of interest in the CACNA1G-AS1 module and their shared miRNAs. b) Log2-transformed and normalized expression values of the three genes of interest from the CACNA1G-AS1 module, TBC1D9, ZBTB16, and MYB, stratified by subtype. c) Log2-transformed and normalized expression values of the three miRNAs targeting the genes in panel b, miR-301a-3p, miR-130b-3p, and miR-301b-3p, stratified by subtype. d) Three genes of interest in the LINC00461 module and of the shared miRNAs. e) Log2-transformed and normalized expression values of the four genes of interest from the LINC00461 module, IGF2BP2, CDK6, RARRES1, and BCL11A, stratified by subtype. f) Log2-transformed and normalized expression values of the miR-190b-5p, which targets the genes in panel b, stratified by subtype.

#### SUPPLEMENTARY MATERIAL

We identified miRNA families and clusters previously reported as important in different cancer types to play a role in the biology of breast cancer. We summarize the main ones here.

a) miR-17~92-1 cluster, mir-106a~363 cluster

Gregorova et al. proposed that the miR-17~92-1 and mir-106a~363 clusters should be considered together since they are frequently upregulated in solid tumors and hematologic malignancies (1,2). We can see a significant visual collection around hsa-miR-18a-5p. hsa-miR-18a-5p in the miR-17~92a-1 cluster that consists of hsa-miR-17, hsa-miR-18a-5p, among other miRNAs (1,2). hsa-miR-18b-5p is part of the mir-106a~363 Cluster. hsa-miR-17 is involved in the ceRNA interactions in three of the 25 modules (ALDH1L1-AS2, ENSG00000230454, LINC00930). hsa-miR-18a-5p and hsa-miR-18b-5p are involved in an significant part of the ceRNA interactions in respectively seven and five out of the 25 modules (ENSG00000261669, ENSG00000232545, ENSG00000249592, ENSG00000240499, LINC02126, (FOXP1-IT1, LEF1-AS1)). hsa-miR-18a-5p and hsa-miR-18b-5p, among other miRNAs of the mir-17~92-1 and mir-106a~363 cluster, were identified as important tumor suppressors in pancreatic ductal adenocarcinoma (3,4). Additionally, decreased levels of hsa-miR-18b-5p were detected in melanoma, inferring the p53 pathway (5) that is crucial in cancer biology (6).

b) miR-130b-3p/301-3p/454-3p family

One of the most significant visual collections in Figure 7 is around hsa-miR-130b-3p and hsa-miR-301b-3p. These miRNAs are as a whole or as a part involved in a large part of the ceRNA interactions in 10 of the 25 modules: ALDH1L1-AS2, TPM1-AS, ENSG00000258535, TMEM26-AS1, ENSG00000229425, BACH1-AS1, LINC00663, ENSG00000225498, CACNA1G-AS1, and DNM3OS. Gregorova et al. described the miR-130-3p/301-3p/454-3p family as significant in cancer biology (2). hsa-miR-130b-3p was confirmed to directly target colony-stimulating factor 1 (CSF-1), downregulate CSF-1 expression, and result in decreased sensitivity to anticancer drugs (7). hsa-miR-130b-3p belongs to the same 22q11.21 miRNA cluster as hsa-miR-301b-3p, which have the same seed sequence and thus share similar target genes (8). Downregulation of this cluster was shown to be inversely correlated with cell proliferation (9), a cancer hallmark (10,11). Furthermore, the cluster was described to play an oncogenic role by Fort et al. (12). hsa-miR-454-3p, the last of this miRBase miRNA family, is not part of the significant collection in Figure 7. hsa-miR-454-3p is described as significant only in the BACH-AS1 module. Decreased levels of hsa-miR-454-3p are associated with less apoptotic cell death and increased cell proliferation (13,14).

c) miR-27a-3p

hsa-miR-27a-3p (19p13.12) is involved in ceRNA interactions in seven out of the 25 modules (ENSG00000259087, CACNA1G-AS1, ENSG00000248671, ENSG00000233178,

ENSG00000261669, ENSG00000232545, ENSG00000249592). hsa-miR-27a-3p can function as both an oncogene or tumor-suppressor, depending on the cellular context (15).

d) miR-34a-5p

hsa-miR-34a-5p (1p36.22) is active in the LINC00461 module and strongly impacts the p53 pathway. has-miR-34a-5p inactivation may substitute for the loss of p53 function in cancer biology (16).

e) miR-190b-5p

hsa-miR-190b-5p (1q21.3) is strongly active in the LINC00461 module. Xie et al. showed in a Kaplan-Meier survival analysis that hsa-miR-190b-5p is significantly correlated with survival in Hepatocellular Carcinoma (17). Inline, Dai et al. suggested that hsa-miR-190b-5p is a potential biomarker for breast cancer overall survival (18). Li et al. showed that high expression of hsa-miR-190b-5p is linked to a good prognosis, while low expression has been shown to drive the malignant progression of pancreatic cancer (19).

#### SUPPLEMENTARY TABLE 1

| | LumA | LumB | Her2 | Basal | Normal | SUM |
|---|---|---|---|---|---|---|
| TCGA-BRCA | 485 | 188 | 74 | 162 | 35 | 944 |
| METABRIC | 679 | 461 | 220 | 199 | 140 | 1699 |

19. Li Y, Wang Z, Zhao F, Zeng J, Yang X. MicroRNA‑190b expression predicts a good prognosis and attenuates the malignant progression of pancreatic cancer by targeting

MEF2C and TCF4. Oncol Rep [Internet]. 2022 Jan;47(1). Available from: http://dx.doi.org/10.3892/or.2021.8223
